# Depression Linked to Frequent Emergency Department Use in Large 10-year Retrospective Analysis of an Integrated Health Care System

**DOI:** 10.1101/115238

**Authors:** Wendy Marie Ingram, Cody Weston, Marylyn D. Ritchie, Sharon Larson

## Abstract

We evaluated general patient features related to depression and frequency of Emergency Department (ED) use in a large integrated health care system. Electronic Health Records of 287,281 adults from a general patient population were studied retrospectively over a 10-year period. Patients with a history of depression were more likely to be seen in the ED and at higher frequency than those without. Frequent ED users were more likely to have a history of depression or psychiatric medication orders than infrequent users. ED visits by depression patients and frequent users have highly correlated complaints and discharge diagnoses with other ED users, often related to pain. Poorly managed depression may be playing a role in frequent ED utilization which may be addressed by universal screening for depression, evaluation of barriers to treatment, and other novel interventions to improve care coordination.

## Introduction

Emergency Departments (ED) are designed to provide patients immediate, unscheduled medical care for acute illnesses and injuries. However, frequent revisits to the ED by a small percentage of patients with chronic conditions or complaints contribute to long wait times and delayed admission of those with critical needs [1–9]. Many studies have implicated depression as a strong predictor of those that present to the ED, especially with complaints of acute or chronic pain and those who present with high frequency [10–17]. In order to address inappropriate overuse of the ED by patients with uncontrolled chronic and psychological conditions, we must better understand the features of these patients and why they continually present to the ED.

To determine these features, we used Electronic Health Records (EHR) from a large integrated health care system in central Pennsylvania to perform a 10-year observational retrospective study of an adult general patient population (population = 287,281). In this descriptive study, we examine patient ED utilization, focusing on those with a history of depression and frequent ED use. To our knowledge, this study is the largest and longest observational EHR-based ED utilization study of a general patient population conducted to date. We discuss our findings and the need for implementation of enhanced protocols for depression screening, increased consideration of depression related care, and evaluation of barriers to treatment, especially for frequent users presenting to the ED with pain complaints.

## Methods

We conducted a retrospective observational study using de-identified Electronic Health Records (EHR) of a general patient population. The study included health care encounter data between January 1^st^, 2005 and September 30^th^, 2015 for patients seen in the Geisinger Health System, an integrated health care system located primarily in central Pennsylvania. Patients 18 years or older at the beginning of the study and who had a Geisinger Primary Care Physician (PCP) during the study period were included in the cohort (n=287,281). Demographic information, medication order histories, and details of ED visits were pulled from a central data warehouse and de-identified by an approved data broker in the Geisinger Phenomics and Clinical Data Analytics Core under the oversight of the Geisinger Internal Review Board as non-human subjects research. Analysis and graphing were conducted with Rstudio (Boston, MA) and GraphPad Prism 6 (La Jolla, CA).

The Geisinger Health System includes six EDs located in central Pennsylvania, one each in Lewisburg, Bloomsburg, Scranton, and Danville, and two located in Wilkes-Barre. The majority of encounters are from the two largest EDs located in Wilkes-Barre and Danville. Patients were defined as having a history of depression if they had an ICD-9 diagnosis code for depression in at least one of four places in their entire EHR, not limited to the study period: 1) an inpatient encounter discharge diagnosis, 2) an ED encounter discharge diagnosis, 3) a problem list diagnosis, or 4) two or more outpatient encounter discharge diagnoses within a two-year period. Patients were defined as frequent ED users if they had four or more visits to a Geisinger ED in a calendar year during the study period [12]. Patient prescriptions for antidepressants, antipsychotics, and antianxiety medications reflect the existence of at least one medication order in a patient's entire EHR, not limited to the study period, and does not necessarily reflect a current prescription at the time of the study nor at presentation to the ED.

## Results

The general patient population included 287, 281 individuals. The cohort is 54% women and 96% white, with 20.6% (n=59,097) meeting the criteria for a history of depression and 79.4% (n=228,184) that do not (Table 1).

**Table 1.**
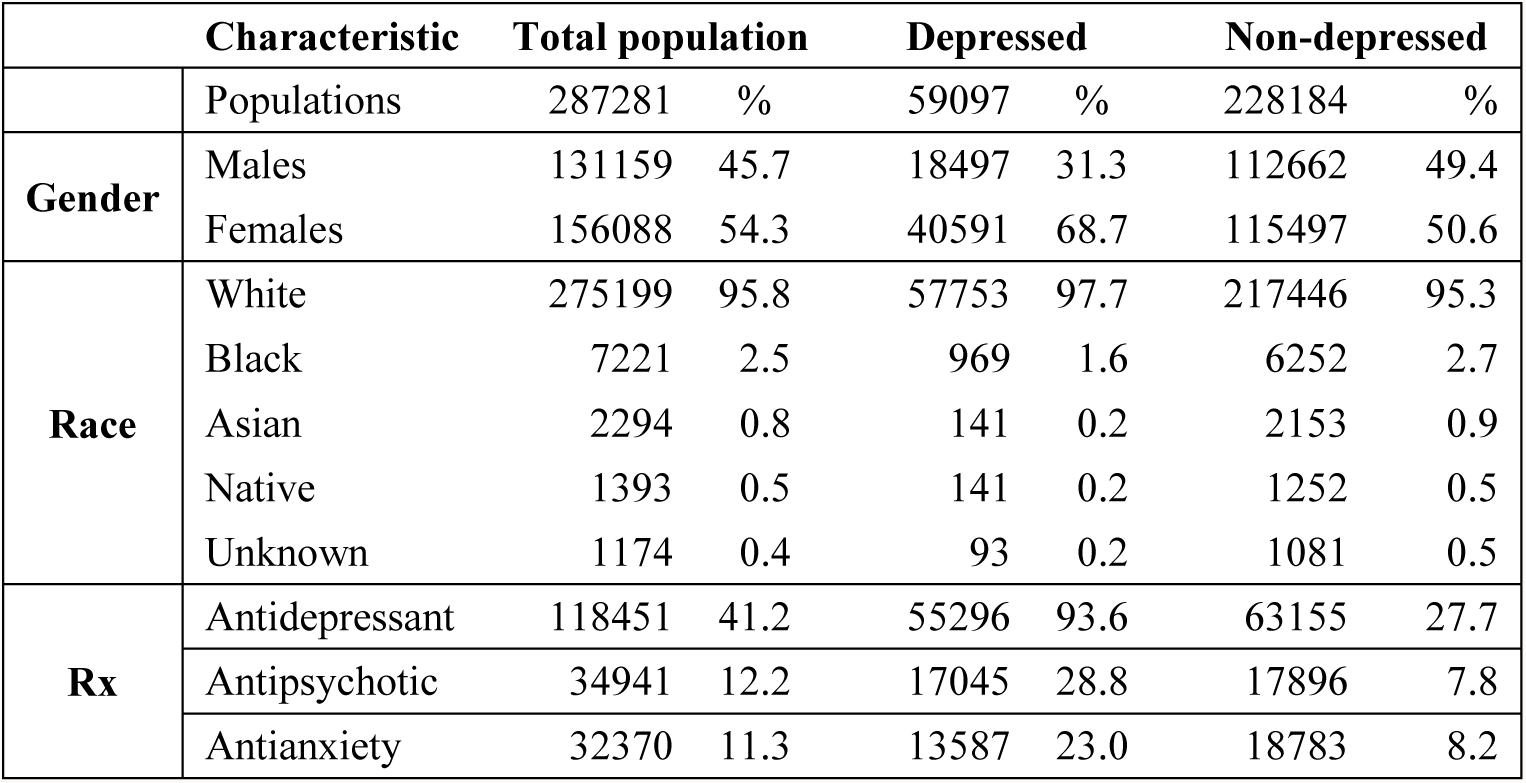
Demographics and medication order history for all patients, those with a history of depression, and those without.

|  | Characteristic | Total population |  | Depressed |  | Non-depressed |  |
| --- | --- | --- | --- | --- | --- | --- | --- |
|  | Populations | 287281 | % | 59097 | % | 228184 | % |
| **Gender** | Males | 131159 | 45.7 | 18497 | 31.3 | 112662 | 49.4 |
|  | Females | 156088 | 54.3 | 40591 | 68.7 | 115497 | 50.6 |
| **Race** | White | 275199 | 95.8 | 57753 | 97.7 | 217446 | 95.3 |
|  | Black | 7221 | 2.5 | 969 | 1.6 | 6252 | 2.7 |
|  | Asian | 2294 | 0.8 | 141 | 0.2 | 2153 | 0.9 |
|  | Native | 1393 | 0.5 | 141 | 0.2 | 1252 | 0.5 |
|  | Unknown | 1174 | 0.4 | 93 | 0.2 | 1081 | 0.5 |
| **Rx** | Antidepressant | 118451 | 41.2 | 55296 | 93.6 | 63155 | 27.7 |
|  | Antipsychotic | 34941 | 12.2 | 17045 | 28.8 | 17896 | 7.8 |
|  | Antianxiety | 32370 | 11.3 | 13587 | 23.0 | 18783 | 8.2 |

### Patients with a history of depression were more likely to be seen in the ED and have a higher frequency of visits than patients without

In the general patient population, 32% of patients had been seen at least once during the study period. Of those with a history of depression, 45% (n=26,605) had been seen in the ED at least once, compared to only 29% (n=65,245) of those without a history of depression (Table 2) (Chi squared test, p = 2.2^−16^).

**Table 2.**
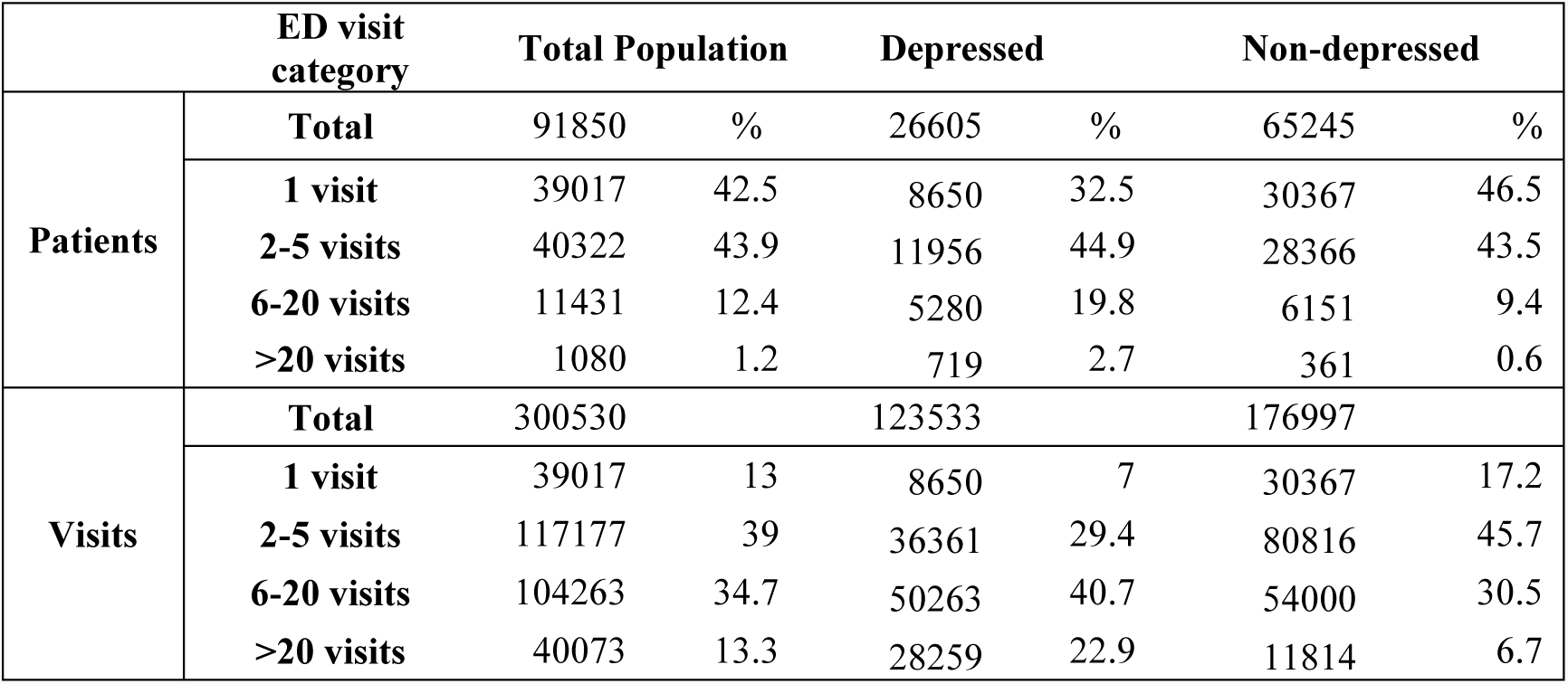
Emergency department use by all patients, those with a history of depression, and those without.

Of the patients seen 6 or more times over the course of the study period, the depressed group had more patients being seen more often (68% of visits by 23% of individuals) than those without depression (37% of visits by 10% of individuals) (Figure 1 A-D). A third of patients with a history of depression and nearly half of those without depression history have been seen only once during the study period (Figure 1 A, B). The higher frequency of visits by patients with a history of depression versus those without (average = 1.8 versus 1.4 visits per patient per year, respectively) is a consistent trend throughout the study period (Figure 1E).

**Figure 1.**
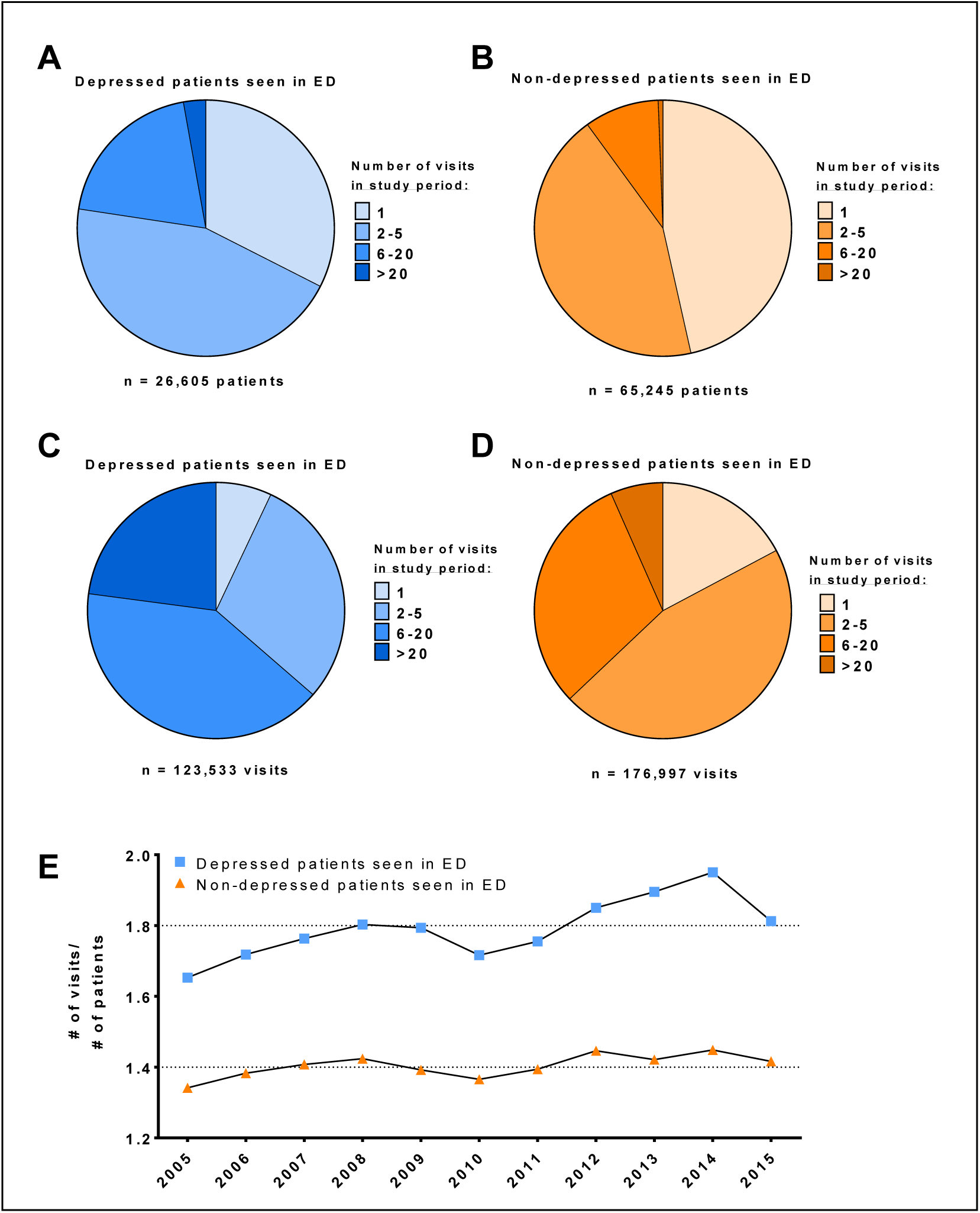
Emergency Department use based on history of depression. A) Percentage of depressed patients (blue), B) percentage of non-depressed patients (orange), C) percentage of depressed patient visits, and D) percentage of non-depressed patient visits for patients that were seen once, two to five times, six to twenty times, or more than twenty times during the course of the entire study period. E) Proportion of visits per patient for each category for each year of the study period.

### Frequent ED users have higher rates of depression history, antidepressant, antipsychotic, and antianxiety medication use than infrequent users

High frequency users are a large and consistent burden on the ED. Of those patients seen in the ED each year, approximately 5% of patients are seen 4 or more times, a consistent trend throughout the study period (Figure 2). These users made up only 2% of the general adult patient population. Those that were defined as being frequent users over the course of the study period consist of 7% (n=6525) of all those seen in the ED and account for 32% (94,386 visits) of all ED visits, averaging 14.5 visits per patient during the entire study period (Table 3).

**Figure 2.**
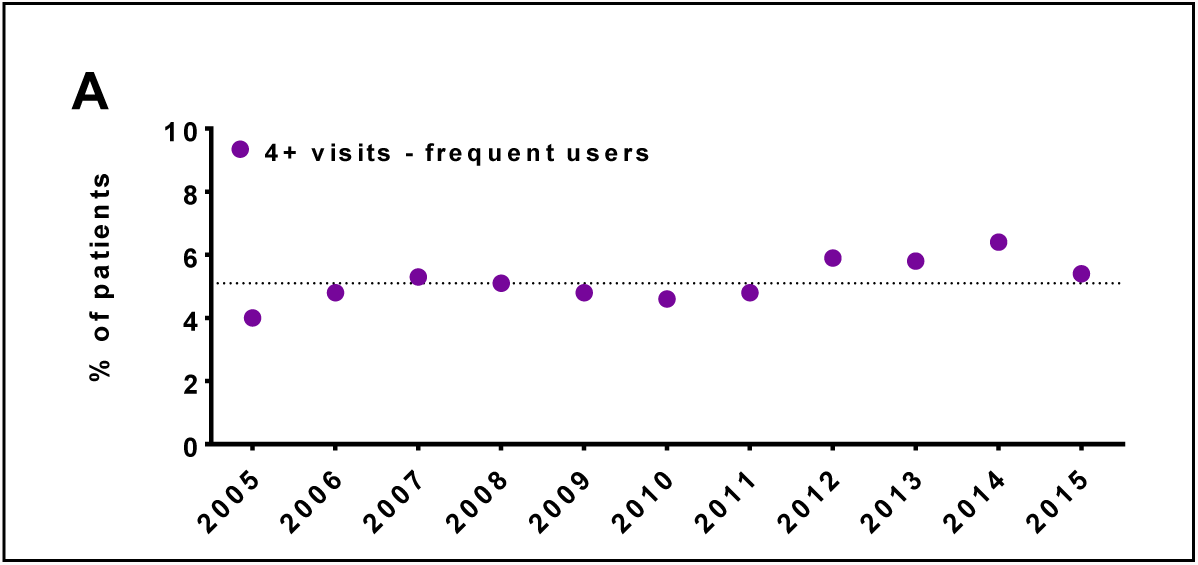
Emergency Department use based on frequency of use. A) Percentage of patients each year that were seen four or more times (purple circles, average = 5.2%) in the indicated calendar year.

**Table 3.** Frequent and infrequent Emergency Department user demographics, depression and medication history.

| Characteristic |  | Frequent users |  | Infrequent users |  |
| --- | --- | --- | --- | --- | --- |
|  |  | 4+ times in 1 year |  | 1-3 times in 1 year |  |
| Patients |  | 6525 | % | 85325 | % |
| Visits |  | 94386 |  | 206144 |  |
| Gender | Males | 2653 | 40.7 | 36942 | 43.3 |
|  | Females | 3870 | 59.3 | 45086 | 52.8 |
| Race | White | 6246 | 95.7 | 82607 | 96.8 |
|  | Black | 230 | 3.5 | 1963 | 2.3 |
|  | Asian | 17 | 0.3 | 360 | 0.4 |
|  | Native | 27 | 0.4 | 250 | 0.3 |
|  | Unknown | 5 | 0.1 | 139 | 0.2 |
| Risk Factors | Depressed | 3216 | 49.3 | 23389 | 27.4 |
|  | Antidepressant | 4638 | 71.1 | 43359 | 50.8 |
|  | Antipsychotic | 3274 | 50.2 | 17701 | 20.7 |
|  | Antianxiety | 2060 | 31.6 | 13226 | 15.5 |

Nearly half of frequent ED users have a history of depression and have high rates of psychiatric medication history (71% - antidepressants, 50% - antipsychotics, 32% - antianxiety). Infrequent ED users have much lower rates of depression and psychiatric medication history (27% - depression, 51% - antidepressants, 21% - antipsychotics, 16% - antianxiety), but these are still higher than the general patient population (20% - depression, 41% antidepressants, 12% - antipsychotics, 11% antianxiety) (Table 3, Table 1).

### The most common reasons for presenting to the ED are primarily for pain related complaints, not mental health concerns

In general, the complaints that depressed patients present to the ED with are similar to those of non-depressed patients (Figure 3). Depression patients have slightly higher frequency of headache- and migraine associated visits, while those without depression are more likely to present with lacerations (Figure 3).

**Figure 3.**
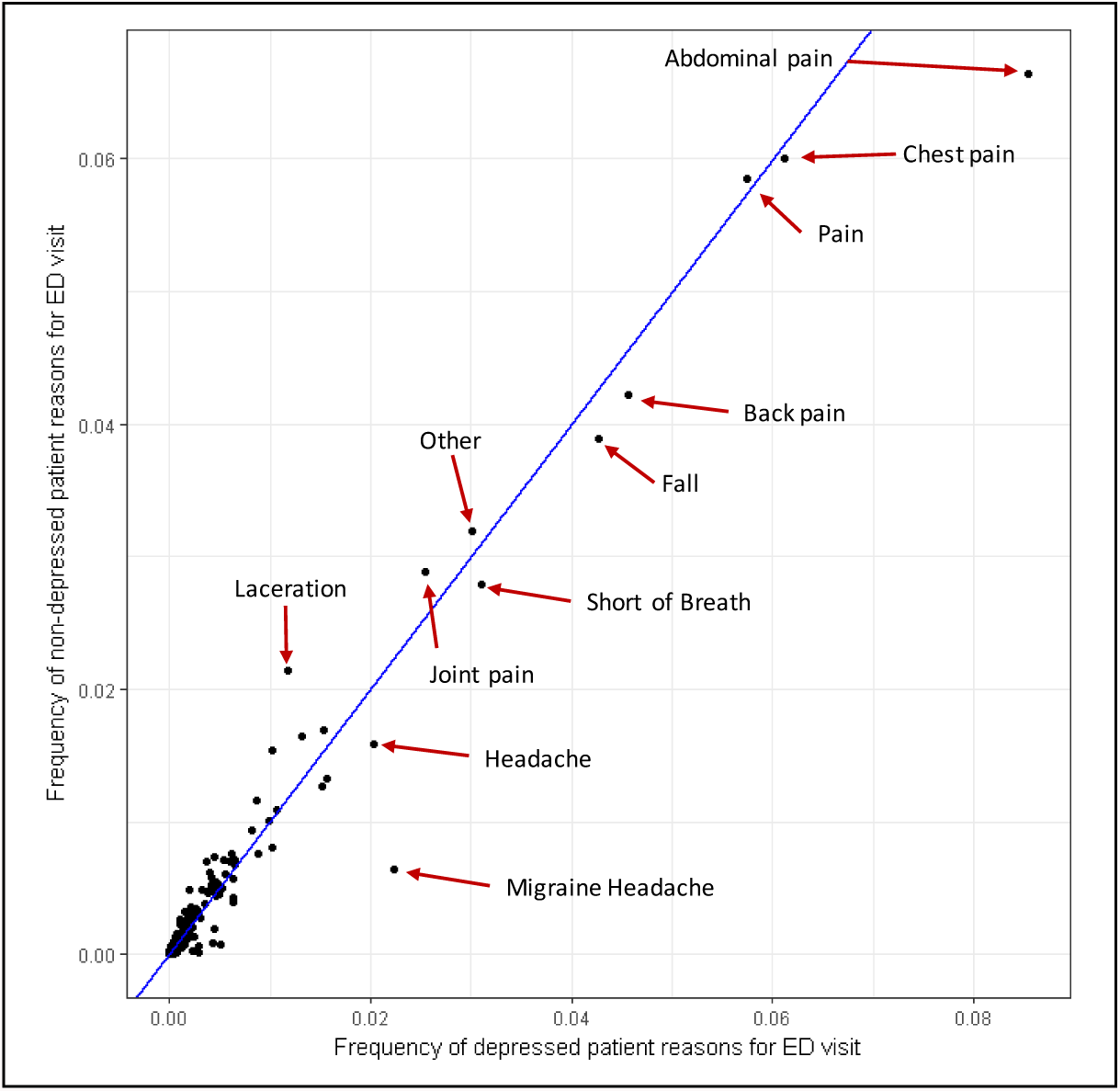
Correlation plot of frequency of reason for ED visit between patients with a history of depression and those without. A x=y perfect correlation line is denoted by the blue line.

Depression patients tend to be discharged at a higher frequency for chest pain, abdominal pain, headache, migraine and depressive disorder, while non-depressed patients are more likely to be discharged with diagnoses of open wounds of the finger, and dizziness and giddiness (Figure 4). Frequent and infrequent ED users are also discharged with largely correlated frequencies of diagnoses (Figure 5). However, frequent users are discharged with codes for abdominal pain, headache, back pain, migraine, and other chronic and acute pain more often than infrequent users. In contrast, infrequent users are more likely to be discharged with diagnoses of chest pain, open wounds of the finger, dizziness and giddiness, and syncope and collapse.

**Figure 4.**
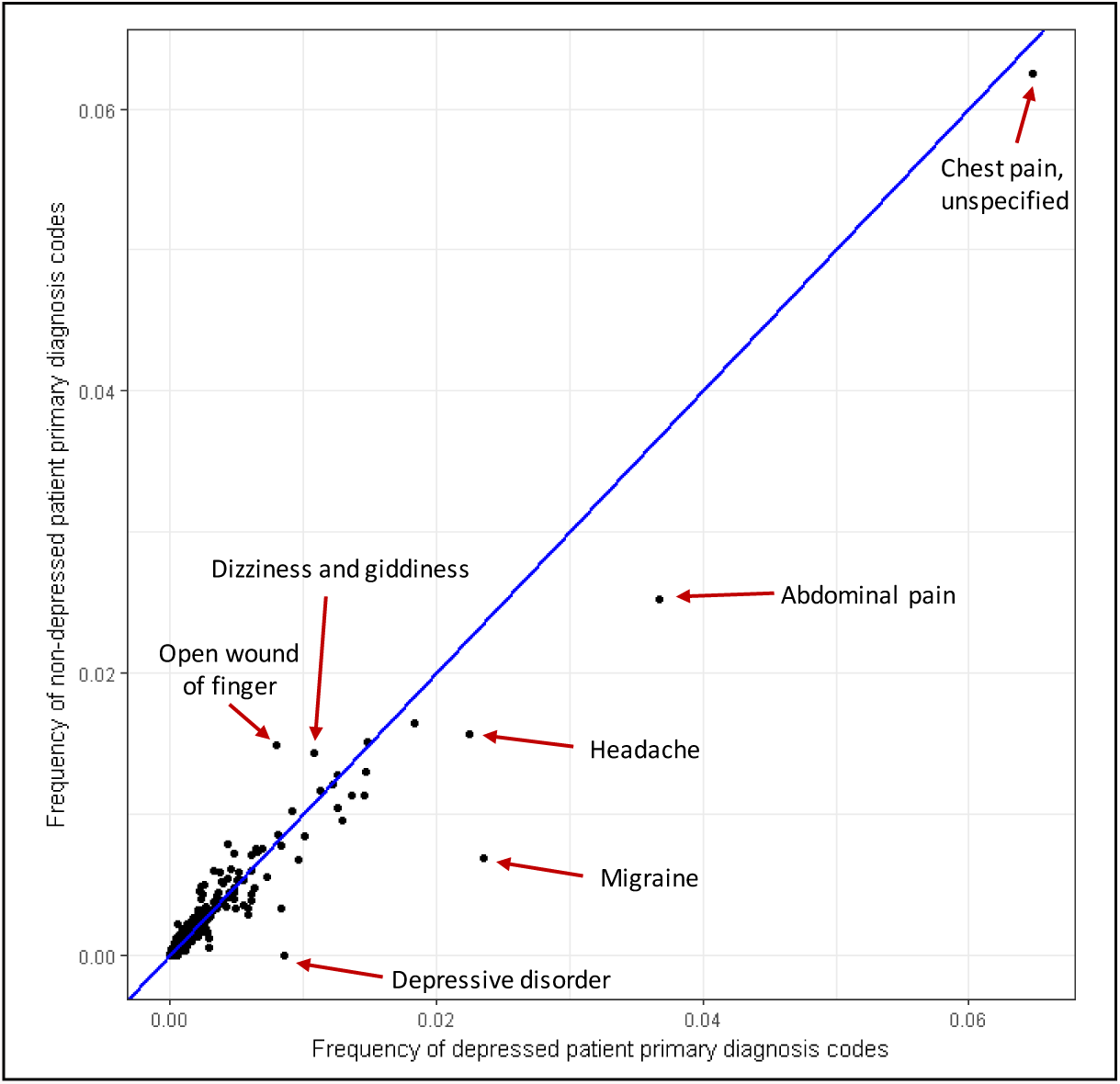
Correlation plot of frequency of ED primary discharge diagnosis between patients with a history of depression and those without. A x=y perfect correlation line is denoted by the blue line.

**Figure 5.**
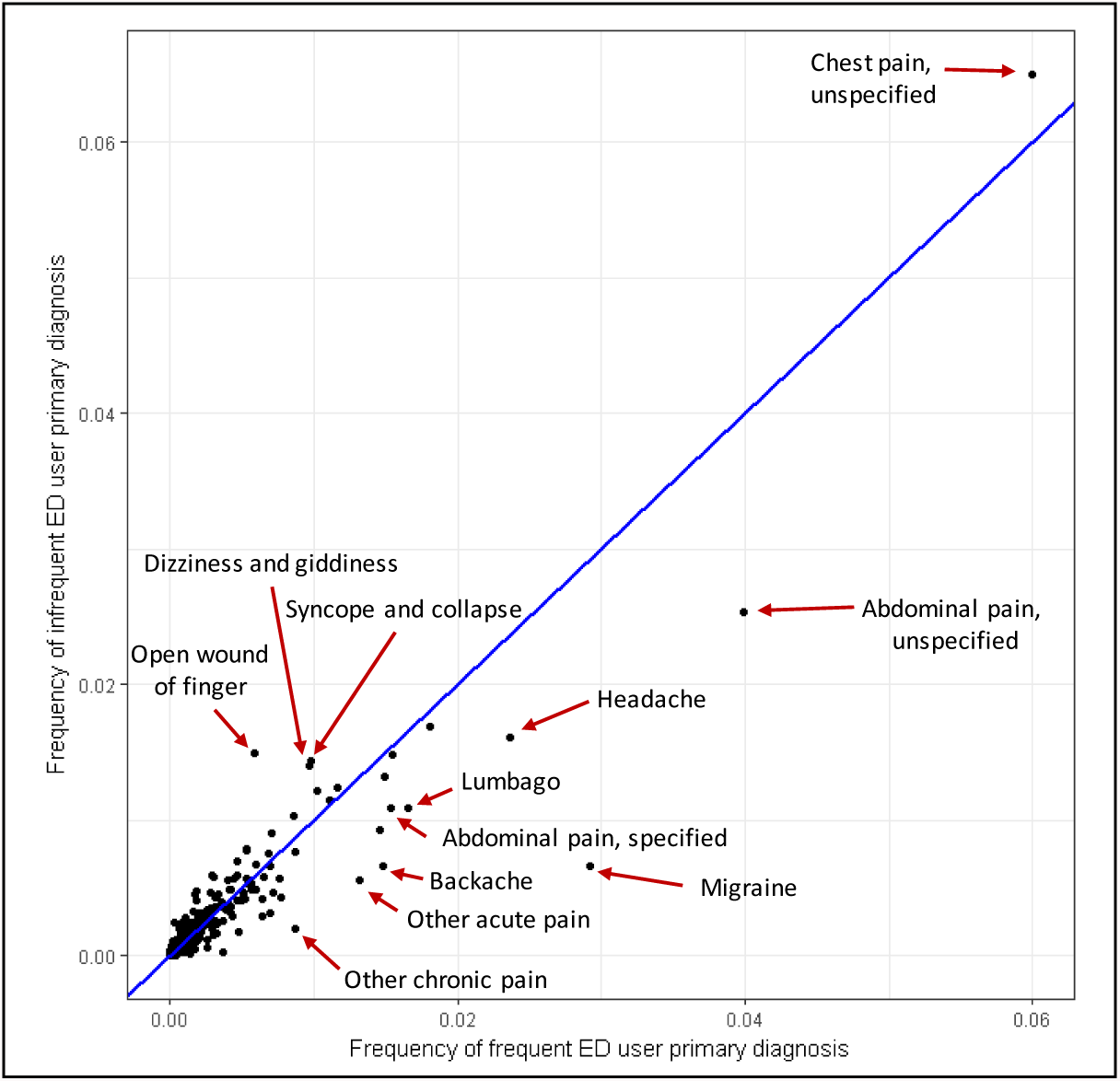
Correlation plot of frequency of primary discharge diagnosis between frequent ED users and infrequent ED users. A x=y perfect correlation line is denoted by the blue line.

## Discussion

In this large retrospective observational study, we have shown that depression and psychiatric medication order history are correlated with frequent non-psychiatric presentation to the ED in a rural integrated hospital system. Depression history is associated with higher rates of ED use, but these patients are not commonly presenting for overtly mental health related complaints. Instead, patients with a history of depression and those that are frequent ED users tend to revisit the ED for various acute or chronic pain conditions. Depression may be highly correlated with frequent nonpsychiatric complaint ED use due to either unrecognized somatic manifestations of depression itself or depression related exacerbations of other comorbid conditions. The interaction between physical symptoms and mental health is difficult to tease apart. In ED settings, both patients and clinicians may be more likely to focus on the physical aspects of disease and overlook the dramatic impact that depression may have on patient symptomology. Taken together, our results support enhanced screening, consideration, and improved management interventions for depression in the ED.

### Depression is related to ED utilization for non-psychiatric complaints

Although the concept of psychosomatic medicine is far from new, we wish to highlight the importance of considering psychiatric factors in improving health care utilization and outcomes. In many cases, patients may be exhibiting somatic manifestations of depression or may be experiencing exacerbation of medical conditions due to a psychiatric illness that is not optimally controlled [6,14,15,18]. Our findings are consistent with previous reports in showing that depression-associated ED use is often seen in the form of various types of chronic pain [3,12,19]. These reports demonstrate that depression is associated with worse prognosis and increased ED usage in abdominal pain, headaches, and general chronic pain conditions. Congruently, our study's highest yield diagnoses are headaches, migraines, and abdominal pain in those with a history of depression. There are complicated biopsychosocial neurofeedback mechanisms underlying the interaction of psychological distress and pain [20–24] which could result in better care and outcomes if considered in the ED setting.

### Somatic manifestations of depression can be misdiagnosed as non-depression related conditions

A wide variety of physiological symptoms, typically considered somatic or “medical” in nature, in addition to psychological or behavioral symptoms, are actually well known symptoms of depression. Depression can result in sleep disturbances, digestive issues and loss of appetite, fatigue and loss of energy, increased pain sensitivity, local or general pain, as well as dizziness [25]. Stress and inflammation are also implicated in causing or exacerbating somatic depression symptoms, either through the activation of the hypothalamic-pituitary-adrenal axis or through inflammatory cytokine signaling [26–28]. Unfortunately, it is not unusual that patients presenting to the ED with complaints of this nature are unaware that these may be related to inadequate treatment for psychiatric conditions. In addition, it is not uncommon for physicians themselves to avoid treatment for their own psychiatric conditions due to stigma [29] which may impact their willingness to consider the physical complaints of patients in the context of mental illness.

### Depression is known to contribute to exacerbations of other comorbidities

Even when there are clear non-psychiatric reasons for presentation to the ED, depression is associated with increased ED use [10]. This has been shown in patients with diabetes, elderly patients, and general ED cohorts [4,30,31]. While major depression's most obvious serious outcome is suicidality, depression has a variety of often overlooked negative consequences which impact the control of patient comorbidities. For example, depression leads to increased medication non-adherence, which can obstruct the medical management of many other illnesses [32]. Diabetes, one of the more well-studied comorbidities of depression, is a striking example of the dangers of undertreated depression. Patients with depression and diabetes use the ED more often than patients with diabetes alone, leading to a 4-fold increase in health care costs [30,33]. More importantly, depression is a negative predictor of survival in diabetic populations, suggesting that the increased use of services and increased costs are not providing enough of a survival benefit to offset the deleterious effects of depression [34]. Distressingly, the prevalence of mental health disorders have increased dramatically in the last seven years [35]. Given the relationship between these conditions and frequent ED use, this does not bode well for future burdens on emergency services [36,37].

### Emergency Department care may benefit from enhanced, depression-focused evaluation and interventions

Medical treatment is often complex, especially in the emergency setting. In the absence of an immediate life-threatening problem, psychiatric concerns can quickly fall to the low end of a long priority list. For example, a study of depression treatment in primary care settings found that medically complex patients often failed to receive care for their depression [3 8]. In ED settings, 50% of patients presenting with tension headaches were found to be depressed, but fewer than one in five with psychiatric need received psychiatric consults [19]. Even when psychiatric problems are recognized in the ED, follow up is often poor [17]. Fortunately, there are many ways to improve quality of care for those with psychiatric complications.

A number of precedents already exist showing that psychiatric intervention in emergency settings can be implemented effectively. Universal screening for suicide risk in the ED increases detection of risk and is widely accepted in both pediatric and adult settings [39–41]. The provision of prescription assistance can help address socioeconomic factors known to increase ED usage [42]. Depression symptom severity correlates with increased barriers to health care access, which contributes to increased rates of ED use [18]. Identification of these barriers and enhanced case management may allow for significant improvements in ongoing treatment, critical to reducing the burden of frequent revisits to the ED.

## Conclusions

Failure to consider depression status and psychiatric history results in suboptimal care, outcomes, and preventable frequent use of the ED. This increases health care costs and reduces efficiency. EDs are often already overburdened, so it stands to reason that appropriate recognition of mental health comorbidities could improve health care quality in this setting. The current standard of care for depression consists of antidepressant medications and outpatient therapy that require consistent patient participation over time. Therefore, even the most psychiatrically-oriented emergency physicians will not be well positioned to provide adequate treatment for this condition. Compounded with the fact that most emergency presentations are not explicitly psychiatric, ED physicians are at a distinct disadvantage in addressing the phenomenon of depression-related frequent ED use. However, recognition that this population of frequent ED users differ in identifiable patterns from the general patient population presents an opportunity to intervene and refer frequent users to more effective treatment resources. In addition, knowing that the proper treatment of depression is likely to improve outcomes for these patients while reducing burden provides a potential quality improvement target for increasing long term population health and efficacious use of emergency resources.

## Acknowledgements

We would like to thank Jason Brown from the Geisinger Phenomics and Clinical Data Analytics Core for his tireless work refining the data pull, and both Christopher Bauer, PhD and Anna Baker, PhD for their excellent comments and suggestions on the manuscript.

